# Towards a Mechanical Model for anisotropic Glioma Spread using Darcy’s law

**DOI:** 10.1101/2021.05.24.445449

**Authors:** Margarita Reséndiz

## Abstract

The growth of a tumor within a finite domain (skull) generates mechanical forces that alter the physical interactions among cells. The relationship between these forces and the tumor architecture remains an open problem subjected to extensive research. Recently, it has been determined that those regions of high mechanical compression can accelerate and intensify the invasive capacity of the malignant cells, forming an irregular tumor whose full extent and edges are difficult to identify.

In the present paper, we propose a one-dimensional mathematical model that describes the process of proliferation and diffusion of glioma cells taking into account the mechanical compression generated during its expansion. Supported on the mixture theory, we model the brain-tumor system as a multiphase mixture of cancer cells, healthy cells, biological fluids and extracellular matrix whose densities determine the mechanical loads generated during the volumetric growth. Our model provides a detailed understanding of the pressure distribution on the interface boundary between healthy and cancer cells. It validates the hypothesis that the conferred ability of cancer cells to proliferate depends strongly on the mechanical pressure sensed. Through the analysis of the mechanical pressure, we determine that the anisotropic loads promote cancer cells to grow preferentially in the directions of low mechanical compression.

## I. INTRODUCTION

The brain is the organ that controls the central nervous system of human beings. It is responsible for monitoring and regulating most of the activities of the body through a sensory system that receives, integrates, and processes information to perform a specific task [1]. The brain is constituted mainly by both neurons and glial cells organized in a highly sophisticated array. The amazing diversity of neurons -whose shapes and sizes are related to their functional role-play the essential role of receiving and transmitting chemical and electrical signals among the brain constituents; while glial cells control the dynamical equilibrium of the brain (i.e., temperature, fluid balance, p.H., blood sugar level), provide support for the neurons and modulate the speed at which the information travels from one nerve cell to another, to mention a few examples. The biochemical and electrical activities are widely accepted and verified as paramount processes that preserve a healthy brain performance. However, they are not the only mechanisms of regulation. Experimental results have revealed that mechanical forces and physical interactions among cells are critical components for regulating the complex dynamic among cells, and play a crucial role in controlling the stress, strain, and pressure to preserve the natural equilibrium of the brain and its normal activity [2–6]. Such mechanical interactions play a central role in safeguarding both brain architecture and brain function [7, 8]. The brain architecture is regulated by anisotropic adhesive forces that preserve its morphology through careful control of its population and a specific balance of adhesive forces. When such balance is perturbed, the mechanoreceptors of each cell sense the changes in the mechanical forces and transform the mechanical stimuli into highly organized electrical signals to recover the biological and mechanical stability [5, 9, 10]. However, it is not always possible to re-establish such equilibrium. Neurons and glial cells are susceptible to multiple and irreparable genetic disorders that affect the regulative mechanisms irreversibly which leads to the loss of their biochemical equilibrium [11–13]. Such genetic disorders are accumulated over the years and frequently expressed by a variety of diseases (A review of diseases derived from glial cell mutations can be found in the reference [14, 15]).

Specifically, if that genetic disorder affects the regulative mechanisms of glial cells, the brain can face one of the harshest complications: a brain tumor. At its more elemental level, the tumor initiation occurs when a mutated cell (or cancer cell) and their daughter cells neglect the biological mechanisms that control proliferation and cell death. The loss of sensibility to the biochemical stimuli trigger an uncontrolled increase of the population due to its rapid proliferation and lack of response to the signals of programmed cell death. At an early stage, the population of cells (≲ 10^7^) are scattered and cannot be condensed in a tumoral mass, so that the tumoral mass is usually imperceptible with clinical devices [16]. The detection of the condensed population in a first clinical diagnosis occurs when the cells are grouped and form an irregular compact mass. At this relatively late stage, each mutated cell has doubled around 25 times and the cells together have reached a total volume of about 1.5 mm of diameter (≳ 10^8^ cells).

Tumor’s classification depends on several factors, such as the kind of mutated cell, the proliferation rate, the speed of spread, and the location of itself in the brain [17–21]. Such details allow the categorization of the tumor from Grade-I to Grade-IV, based on abnormalities of the individual cells. Grade-I tumors are often benign and not diffusive, but can irreversibly evolve toward higher grades; while Grade-II, III, and IV are malignant tumors whose aggressiveness get worse as their grade increase. Each category represents undoubtedly a risk to the patient, but a Grade-IV tumor –known as *Glioblastoma Multiforme* (GBM) or glioma– has a remarkable aggressiveness due to its ability to invade and migrate quickly towards the track of the white matter [22]. These characteristics of rapid proliferation and invasion are the main lethal factors that prevent precise treatment and diagnosis in patients, causing quite a low probability of survival. The rapid proliferation and vascularization of GBM inside the brain tissue make the tumor mass indistinguishable from the healthy tissue so that root part of the tumors, the part that invades the healthy tissue, can never be defined accurately under clinical imaging devices, making its treatment even more difficult. In other words, the low density regions of cancer cells are undetected or confused with healthy tissue as a consequence of the lack of resolution in medical devices.

Glioma cells, on the other hand, are characterized for being multiform cells whose nuclei are oval or elongated, and with a bigger size than the healthy cells [23]. Despite its apparent morphological differences between healthy and cancerous tissue, such criterion cannot be used to specify the location of a tumor in vivo. They display significant variation in growth rates among patients so that it is unfeasible to determine a global growth rate that helps to predict its growth in the individual patients [24, 25]. Through the statistical analysis, it has been determined that the median specific growth rate of the tumors is 1.4% per day, and the equivalent volume doubling time was 49.6 days. These results provide a general scheme but cannot be used to identify/predict the full extent in individual cases.

The dynamic of growth of a tumor through mathematical modeling has been a complementary paramount tool to understand some of the complex processes that take place during the spatio-temporal dynamic of glioma cells, and predict the full extent of the tumor. Some of the investigations with significant influence on explaining the progression of the tumor at its more elemental level are reported in the references [21, 24, 26–30]. In these studies, the gradual growth of a tumor is described through a law of exponential growth to represent the proliferation process, and a Gaussian function imitating the infiltration of glioma cells in the healthy tissue. The obtained results by such models reproduce the global architecture of the tumor satisfactorily but also has differences with realistic situations. In this sense, new proposals have arisen not only to identify and understand the components that control the growth, invasion, and migration of cancer cells but also to investigate their geometries, with the aim of improving the estimation of the full extent of the tumor [21, 24, 26–30]. Murray, and Swanson et al., for instance, model the preference of glioma cells to diffuse through white matter over gray matter [21, 24]. Woodard et al., improve such a proposal using the specific anatomical structure of the brain obtained from Computerized Tomography (CT) and Magnetic Resonance Imaging (MRI) of a patient [26]. Chaplain and Sleeman’s study model the chemotactic agents that promote the cancer cell diffusion toward the brain tissue and the influence of the chemical stimulus on the kinetics of tumor shape [31], Macklin, and Lowengrub consider the genetic characteristics of the tumor as well as the microenvironment surrounding it [32].

All these models and many other scientific papers, have provided different and relevant contributions to understanding the physics behind the tumor growth, so that the precise delimitation of the tumor shape has substantially improved. Nevertheless, despite abundant research, there is scarce knowledge about the biochemical and physiological impact of the mechanical forces on the brain tumor, there are several open questions and the precise characterization of the full extent is still poor. Therefore the tumor growth mechanisms require a formal description using basic fundamentals and a multidisciplinary analysis.

In the present paper, we construct a bio-mathematical model that describes the temporal-space evolution of glioma cells of a solid avascular tumor growing in a finite domain. The aim of this paper is to improve the characterization of the shape of the tumor by adding a new consideration; the mechanisms of expansion and compression in the processes of diffussion and proliferation.

In order to present this argument, in section II, we make a review of the mechanical implications of the tumor growth at mesoscopic and macroscopic scales. In section III, we present the standard mathematical construction of the brain tumor model, which will employ Darcy’s Law, this time proposing a novel manner to express quantitatively the transportation of cancer cells via mechanical compression. The one-dimensional representation is displayed in section IV, where we analyze the isotropic and anisotropic growth. Finally, in section V, we discuss our results.

## II. MECHANICAL ASPECTS OF TUMOR GROWTH

At a cellular level, the processes of proliferation and diffusion of glioma cells in a finite domain (the skull) are accompanied by gradual deformations in both the glial and healthy cells [3, 28]. The increased population of the tumor due to the proliferation, compresses and deforms the architecture of the neighboring cells and consequently, modifies the mechanical properties among them such as pressure, adhesive forces, stress, and strain [3]. Recent studies have established that alteration of mechanical forces in a healthy tissue arise out of the demand for specific biological functions [33, 34]. Nevertheless, altering the mechanical forces of tumor cells can benefit and intensify the invasion and metastatic capacity of cancer cells [4].

Currently, there is no logical connection between the morphology of a cell, its biological role, and the effects of the mechanical loads, so that the topic is an active area of research. Nevertheless, it is known that morphological changes derived from mechanical alterations on gliomas increases their stiffness. This characteristic, probably the only one visible on tumors, trigger the production of the extracellular matrix and stromal tissue, increasing the rigidity of the malignan cells [35–38]. Additional reports suggest that the local tissue stiffness promotes the tumor growth, activating the proliferation pathways of malignant cells, and has the potential to alter both the function and activity of glial cells [35]. At a macroscopic level, on the other hand, the growth of the malignant mass in a restricted space induces a system of forces (loads) between the solid tumor and the healthy tissue. The tumor expansion exerts an outward force that compresses, displaces, and deforms the healthy tissue. In contrast, the healthy tissue exerts an opposite inward force on the tumoral mass in order to compete for the physical space and avoid being compressed. The architectonic changes derived from these mechanical forces are known as *mass effect*. In a more detailed analysis, it has been identified that the mechanical loads generated during the volumetric growth produce total solid stress as a combination of i) externally applied stress and ii) growth-induced stress [39, 40]. The externally applied solid stress is the physical force exerted by the tissue on the tumor. The physical force that compresses the tumor during its volumetric expansion is known as externally applied stress and it effects the tumor differently. In the internal region, the forces are directed inward radially in a way that compresses the cells into the tumor center, while on the tumor’s surface there are two kinds of force vectors: a compressive force that tends to compact the cells toward the center of the tumor that presumably, can either inhibits the cellular division and triggers apoptotic cell death or enhance the invasive phenotype of cancer cells and increase the expression of genes that remodel the ECM and tumor vessel [40], and a tensile force that is tangent to its surface. The growth-induced solid stress, on the other hand, is the physical pressure accumulated within the tumor as the cancer cells proliferate. The mechanical force generates strain energy that pushes and deforms the brain tissue and strains the tumor microenvironment. Hypothetically, such forces regulate the growth rate and apoptosis [40].

Because the heterogeneous structure of the brain and the tumoral mass have different infiltration levels on white matter and anisotropic diffusion rates, the forces produced are also inhomogeneous and anisotropic. Thus each part of the tumor is exposed to various mechanical loads and consequently possesses different physical properties described by different mechanical behaviors. A strong hypothesis suggests that such anisotropic mechanical loads can induce apoptosis in regions of high compressive stress and allow a proliferation of the low-stress areas of the tumor spheroids [4]. Anisotropic loads force cancer cells to grow preferentially in the directions of low stress and can shape tumor morphology [40].

### A. Previous modelling of the mass effect

The mass-effect on tumor growth has been mainly modeled using two different perspectives: i) analyzing the stress, strain, and deformations produced by a solid tumor with nonlinear viscoelastic behavior [41–44] and, ii) analyzing the physical interactions among a multi-component tumor constituted by deformable sphere-like structures residing in biological fluids [45–48]. The second scheme’s analysis of the brain tumor has been described at different levels of complexity, ranging from mixtures from two to multiple species [45–48]. These models are based on the principle that all the components co-exist simultaneously, and evolve according to balance laws and conservation principles. Some of the earliest investigations about the influence of the mechanical interactions on solid tumors were performed by Burton, Greenspan, and Thomlinson [49–51]. Their models conceive the lesion as a set of deformable sphere-like structures that proliferate and displace over isotropic space so that each cell is affected by isotropic forces, and the resultant tumor shape is symmetric. Subsequent studies performed by Greenspan improve such a model by considering the asymmetric space where the tumor growth, the nonuniform intercellular forces, as well as the population changes caused by the death or birth of cells [52]. From here, Greenspan suggests that the shape and structure of the tumor are affected by the internal pressure differential produced by the birth or death of cancer cells. In a more detailed description, Macklin and Lowengrub investigate the tumor morphology as a function of the concentration of the surrounding nutrients and determine how such shape is affected when mechanical factors are taken into account [32]. Clatz et al. [53], simulate the glioblastoma growth on a patient-specific model, involving the physical deformation of the brain, derived from the physical forces generated during growth and infiltration processes, and the internal structure of the white matter. Their research models the mass effect taking into account the mechanical contributions originated by two different processes: volumetric expansion and the infiltration (diffusion).

Ambrosi and Mollica model the effects of the mechanical stress generated by the volumetric growth of a multicell spheroid -emulating an avascular tumor- on two growth conditions. The model is adapted to a wide range of geometries and different boundary conditions [48]. Chaplain et al. [31] study the disorders caused by the stress produced by the tumor cell proliferation. The model reproduces some phenomenological states associated with the volumetric growth, such as the progressive compression within the tumor itself, and the compression caused to the surrounding tissue. A more sophisticated model is proposed by Preziosi et al. [45], who represent the brain-tumor system as a mixture of solid components embedded in a liquid media interacting between them. The analysis of the mass-effect is performed through the mechanical interactions among a set of multiphase spheroids representing the different kind of cells, all embedded in a fluid media. The spatial tumor evolution is studied, assuming that the deformation of the tumor mass is correlated with the velocity displacement of the cancer cells toward the healthy tissue.

The implementation of the spatial inhomogeneities of the tissue, as well as the real specific parameters (*e.g.* net proliferation rate), have improved the prediction of the physical boundaries of the tumoral mass. Supported upon the multiple scientific evidence of the mass-effect during tumor growth, we present a mathematical model to explore the shape of a tumor in 1D whose cells not only diffuse and proliferate, but also are exposed to expansive and compressive inhomogeneous mechanical loads in an anisotropic finite space. Appealing to the principles of the theory of mixtures used in the models of Preziosi et al. [45], the brain-tumor system is considered as a multiphase mixture represented by deformable sphere structures, embedded in a liquid phase which contains the extracellular material surrounding the cells. The model is based on the principle that all the components co-exist simultaneously and evolve according to balance laws and conservation principles. Two key elements in our formulation are that glioma cells move in an anisotropic medium obtained from a CT, and the simulation is performed with clinical parameters from patients. For simplicity, we analyze the numerical results in one-dimension with the aim of predicting the direction and extent of the spread of a tumor in an anisotropic medium.

## III. MATHEMATICAL MODEL

We consider the brain-tumor system as a material body constituted by four-components: (i) glioma cells, (ii) healthy cells, (iii) biological fluids and (iv) the extracellular matrix (ECM) [31, 45, 46]. At an early stage, the tumor is conceived as a small cluster of glioma cells embedded in a complex array of components all of them confined within the skull. Each cancer and healthy cell is surrounded by a fluid film (and ECM molecules) that discards dissipative forces as malignant cells diffuse, but preserves the adhesive forces. Referring to the multiphase models proposed in [45–47, 54, 55], the healthy and cancer cells are represented by a set of two different volume fractions. The group with the minority population in the initial stage (gliomas) has the ability to multiply and move inside the wet porous structure, while the other ones (healthy cells) are restricted to limited displacement or no displacement (ECM). The biological fluid is used to balance the other constituents.

Let’s consider an arbitrary control volume 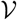 with surface 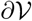 and total volume *V_T_*. The cancer cells, healthy cell logical fluid, and extra cellular matrix residing in 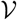, occupy a volume fraction *ϕ_c_*(**x***, t*), *ϕ_h_*(**x***, t*), *ϕ_e_*(**x***, t*), and *ϕ_l_*(**x***, t*) ∈ [0, 1], respectively. In the model, we assume that the mixture is saturated in such a way that the whole space is filled with the four components, and all together satisfy the mathematical constriction ∑*_i_ ϕ_i_* = 1.

Let *ρ_i_* (for *i* = *c, h, e, l*) be the constant mass density of constituent *i* in 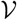. The total mass *M_i_*(**x***, t*) at time *t* = *t*_0_ occupied by the *i*-th constituent in 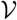 is defined as

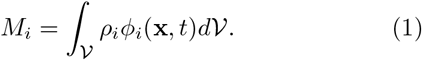

As the tumor increases, the total mass *M_i_* in 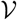 changes due to cell proliferation, cell decay, diffusion or or advection.

Following the principle of mass conservation, the equations that describe the temporal evolution of each constituent *i* − *th* of the mixture in 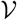, are

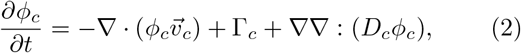

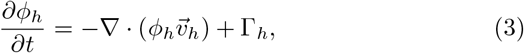

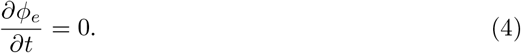

Note that *ϕ_l_* is not modeled explicitly, but it is present via the mathematical relation *ϕ_l_* = 1 − *ϕ_c_ − ϕ_h_ − ϕ_e_*, in order to guarantee the balance of the equations.

In the set of equations, 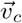 and 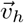 are the velocity fields with which cancer and healthy cells move about in the brain, while Γ_*c*_, and Γ_*h*_ are the growth profile of the cancer cell population defined by the logistic growth law, Γ_*c*_ = *r_c_ϕ_c_*(1 − (*ϕ_c_* + *ϕ_h_*)), and the decay profile expressed as Γ_*h*_ = −*r_h_ϕ_c_ϕ_h_*, respectively. *D_c_*(*x*), on the other hand, is a spatially dependent, anisotropic diffusion tensor that stores the directional anisotropy of the spread of cancer cells at each spatial point along the white matter fibre tracks. As shown in Swan et al. [56] and Painter and Hillen [57] the diffusion tensor *D_c_* can be parametrized from clinical MRI-DTI data (diffusion tensor imaging). In this formulation it shows that the appropriate form of the anisotropic term is not a typical Fickian-kind of diffusion ∇·(*D_c_*∇*ϕ_c_*), but rather the *fully anisotropic*, or Fokker-Planck form: ∇∇: (*D_c_ϕ_c_*) (see [56, 57]).

### A. Constitutive equations derived from mechanical effects

Describing the evolution of a solid tumor using mixture theory requires closure relations for the velocity fields involved in Eqs. (2) and (3). The standard procedure in most formulations requires the momentum balance equations generated from the physical interactions among cells, and the associated constitutive laws [45–47]. Such a procedure is quite promising because it not only considers the infiltration and diffusion of cancer cells into the healthy tissue but also the impact that mechanical compression has on the tumor shape during its progress. However, the model is limited to numerical simulations because of the many unknown biological parameters involved. Our model simplifies the problem by proposing a reduced closure relation that correlates the mass increase in the tumor with the mechanical compression that it generates on itself and the surrounding tissues. First, we start by assuming that the natural configuration of the brain (or zero stress configuration) is associated with a homogeneously distributed mechanical compression (pressure), as depicted in Fig. 1 a). When the proliferation of glioma cells occurs, the increased volume exerts a mechanical force radially outward that gradually compresses the healthy tissue. When the healthy tissue perceives the excess pressure, a combination of mechanical forces between the tumor and the host tissue is produced to induce structural changes in both the tumor shape and host tissue. For simplicity, such pressure differential is considered as a unique consequence of the increased volume, putting aside any changes caused by stress and strain.

**Figure 1.**
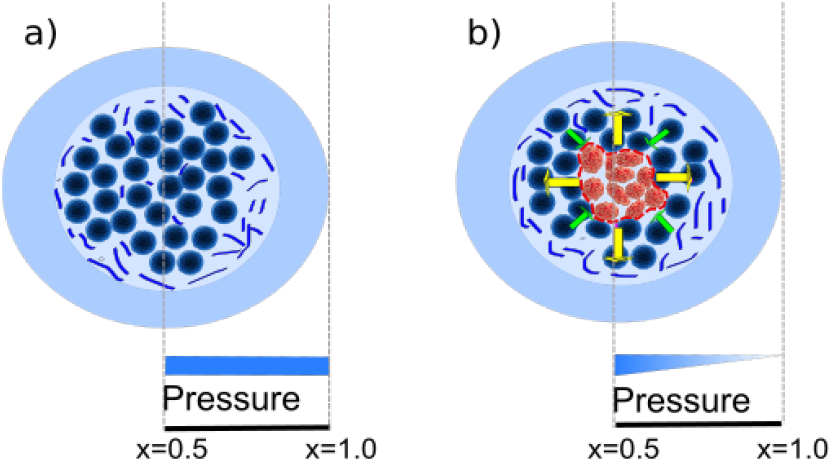
Schematic representation of the mechanical pressure in a brain slice. a) The healthy configuration is associated with a homogeneously distributed mechanical pressure as part of its normal activity. b) When a tumor is present, the mechanical loads generated during the volumetric growth produce a total solid stress with two components: i) externally applied stress and ii) growth-induced stress.

Appealing to physical arguments, the magnitude of the mechanical compression exerted on the tumor cells depends on the population of healthy cells surrounding them. The larger the population of cells surrounding a cluster of cancer cells, the greater the force it exerts on healthy cells to make room for itself. Consequently, the cells located in the center of the tumor exert greater force to make room for itself than those located close to the interphase [46]. Thus, we can say that the maximum pressure occurs in the center of the tumor and gradually decreases toward the interphase among boundaries [55]. Additionally, we consider that such pressure distribution is not isotropic. We take into account that some cancer cells are biologically conferred with nutrients that allow them rapid proliferation with respect to others. This difference assigns an anisotropic cancer cell distribution, and consequently, an anisotropic mechanical compression.

To characterize the mechanical compression, we take into account two basic mechanisms in the finite domain: i) as the glioma cells population increases over time, the tumoral mass increases its volume. Consequently, it enlarges outward, producing a radial compression that lies on the host tissue. ii) When the host tissue perceives the excess pressure, it exerts a repressive force upon the growing mass competing thus for space, as depicted in Fig. 1b).

Following the reasoning of Wasserman et al. [58], the mechanical pressure exerted on a cancer cell can be characterized as a function of the cell density. The mathematical function that quantifies the pressure has the form:

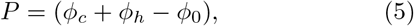

where *ϕ*_0_(*x*) is the density of the host tissue in the zero-stress configuration.

A direct consequence of the mechanical alteration in the tumor morphology is the generation of a drift. The outward pressure produced by the increased volume of cancer cells displaces the cancer cells and healthy cells with velocities 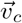 and 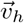, respectively. Additional mechanisms that contribute to the cell drift velocity will be neglected.

The simplest relation between the velocity field and the mechanical pressure is known as Darcy’s Law, which correlates the displacement velocity field of the cells 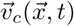, with the mechanical compression derived from the tumor growth. In this simplified model, we consider that the velocity produced is a direct consequence of the mechanical expansion caused by the volume increase and the diffusion of glioma cells. Thus, the relationship of *v* with the pressure *P* is through the expression

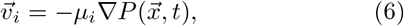

where *μ_i_* represents the ratio of permeability coefficient and viscosity in the medium. Assuming that the displacements are a consequence of physical forces produced during the expansion and that in a multiphase model the velocity field of the cells is the same for each constituent proposed by Ward and King [59–61], the velocities for cancer and healthy cells are proportional, i.e. 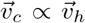. In spite of the fact that we are assuming the components with the same velocity, we will include a factor *μ_i_* to differentiate the ability for the tissue to respond to the pressure differently, i.e.

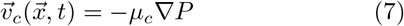

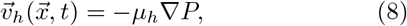

where we impose that *μ_c_ > μ_h_* to highlight that cancer cells have less adhesion and, consequently more movement so that they infiltrate the healthy tissue more easily than does the healthy tissue into the tumor.

In summary, the set of equations (2)-(5) together with the expression Γ_*c*_, Γ_*h*_, (5), (7), and (8) form a closed equation system that describes the tumor growth considering simultaneously the anisotropic diffusion and mechanical compression (mass-effect). In the next section, we will analyze the distribution of cancer cells in a one-dimensional case of a virtual brain tumor modeled with and without the mass-effect. We will study the mass-effect together with the anisotropy on the tumor shape and the an artificial DTI data set.

## IV. NUMERICAL SIMULATIONS IN 1-DIMENSION

In order to study the impact of the mechanical compression on the diffusion of cancer cells in a heterogeneous medium, we explore the numerical solution of the equations (2), (3), and (4) in 1-dimension, assuming *ϕ_e_* constant. The velocity terms defined in (7) and (8), together with the transport equations in 1D, lead to the equations

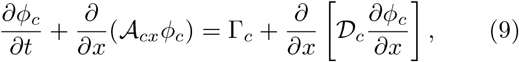

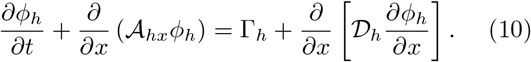

where the effective drift and effective diffusion terms are defined, respectively, in the form

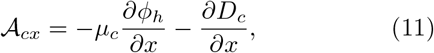

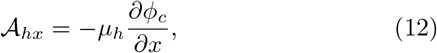

while the effective diffusion coefficients are

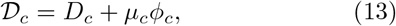

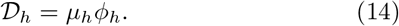

Notice that 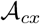 and 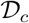 are constituted by the advective and diffusive contributions, respectively, of both − ∇ · 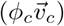 and ∇∇: (*D_c_ϕ_c_*).

The analysis of the temporal evolution of the tumor in 1-dimension is performed through the numerical solution of the coupled parabolic Eqs. (9) and (10). The simulation starts by assuming that in the early stage, the initial population of cancer cells *ϕ_c_*(*x*_0_*, t*_0_) is represented by a Gaussian distribution with variance *σ*, situated around the spatial coordinate *a*. The healthy tissue (or complementary tissue) surrounding the tumor is represented by *ϕ_h_*(*x*_0_*, t*_0_). Both populations are mathematically represented by

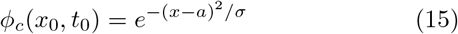

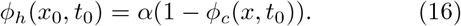

The *α*-factor in Eq. (16) indicates the fraction of extracellular matrix *ϕ_e_* and biological fluid *ϕ_l_* in the total mixture, both of which remain constant in time.

The tumor progression with mass-effect will be studied in three different cases: In subsection A, we analyze the effects of the mechanical compression on a tumor isotropically growing with a constant diffusion coefficient. In subsection B, we introduce the spatial anisotropy using a linear diffusion coefficient, and in subsection C we asses such anisotropic grow using an heterogeneous distribution from an CT of a brain.

### A. Isotropic tumor growth with and without mass-effect

In our model, the spatio-temporal progression of an avascular virtual tumor at an early stage, centered at *x*=0.5, is represented by a Gaussian distribution with variance *σ* ≈ 0.2. This distribution represents the configuration and location of the primary glioma cells that give rise to the tumor lesion. For simplicity, we work in a scaled space of domain [0,1], whose distribution function is restricted to the maximum value of 1. Because we are modeling a real high grade tumor, we assume that the glioma cells have a proliferation rate *r_c_*=0.012/day and a diffusion coefficient 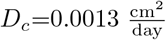 that represents the isotropy of brain [21, 24, 27, 29], as depicted in Fig. 2 (dotted line).

**Figure 2.**
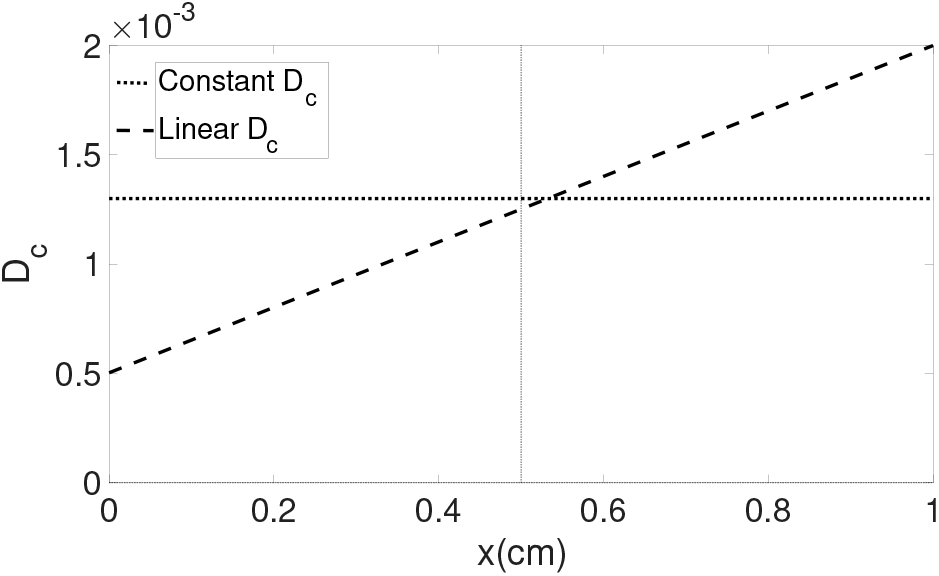
Numerical representation of the diffusion coefficients for isotropy and anisotropy spaces. The isotropy in brain is represented by a constant diffusion coefficient whose value is *D_c_*=0.0013 for a high grade tumor (dotted line). The anisotropy is represented by a linear diffusion coefficient where 0.005< *D_c_* <0.002 for a real high grade tumor (dashed line).

We consider that the decay rate of healthy cells is *r_h_*=0.09/day, and the values of the permeability coefficient are *μ_c_*=0.01 cm/s and *μ_h_*=0.001 cm/s. These magnitudes indicate the rates at which glioma cells invade the healthy tissue, and the healthy tissue infiltrates the tumoral mass, respectively.

We examine the virtual tumor via the numerical solution to Eqs. (9) and (10), at time *t* ≈ 8.7 months after its onset, as depicted in the Fig. 3a). The image displays the profile of concentration of cancer cells (black-solid line) and healthy cells (black-dashed line) undergoing proliferation, diffusion, and mass-effect during the spatio-temporal progression.

**Figure 3.**
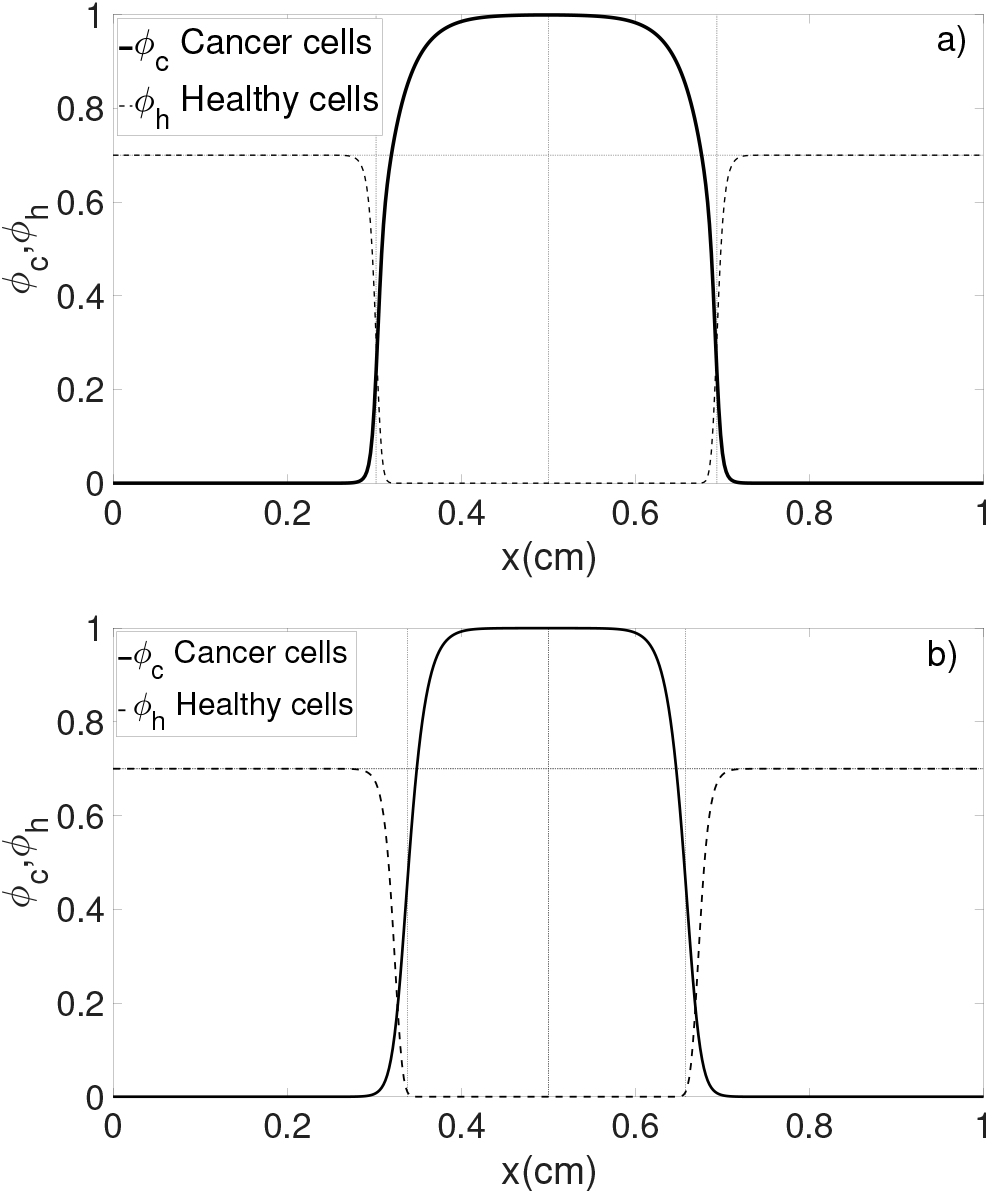
One-dimensional schematic representation of the distribution of cancer cells of a virtual brain tumor modeled with and without the mass-effect. The cancer cell distribution profile is measured at *t* = 8.7 months later, performed in the same conditions with *D_x_* = 0.0013, *r_c_* = 0.012, *μ_c_* = 0.01, and *μ_h_* = 0.001. In a) the cancer cell and healthy cell distribution is displayed with mass-effect and in b) the Cancer and healthy cell distribution without mass-effect.

From the image, we identify that the concentration of cancer cells (black-solid line) is highest around the spatial point where the tumor is originated (*x*=0.5), and decays outward reaching their lowest values once they have infiltrated the healthy tissue. At the same time, the healthy tissue (black line) is destroyed and displaced from the center of the tumor radially outward as the tumor grows. The profile of concentration of both cancer and healthy cells is symmetric around the initial spatial point due to the constant and isotropic diffusion coefficient.

To evaluate the mass-effect on tumor progression, we contrast the growth profile of our model, Fig. 3a), with a purely diffusive-proliferative model (without mass-effect) proposed by Swanson et al. [30]) in Fig. 3b). Both numerical simulations are performed using the same initial conditions, parameters, and time of progression in a rescaled space [0, 1]. The cancer cell density profile of the diffusive-proliferative model has a similar behavior to the model involving mass-effect: the concentration of cancer cells is maximum around the tumor nucleus, and decays as it extends and invades the healthy tissue, reaching the null value once it has infiltrated the tissue. In spite of the similar profiles of both cancer cell distributions, there are slight differences between their curves. The inclusion of the mass effect in Fig 2a) shows a steeper gradient on the tumor boundary as compared to the no-mass effect in Figure 2b). Also the plateau is more flat in the no-mass effect case. The inclusion of a mass effect in this case has a notable, but not dramatic effect.

The influence of the permeability coefficient on the tumor shape is explored in Fig. 4a) for three different values: *μ_c_*=0.01 cm/s, *μ_c_*=0.02 cm/s, and *μ_c_*=0.03 cm/s. As to be expected, the tumor extension (black-solid line) becomes larger as the permeability coefficient increases. Moreover, we see larger mixing of healthy and cancer tissue for increased permeability.

**Figure 4.**
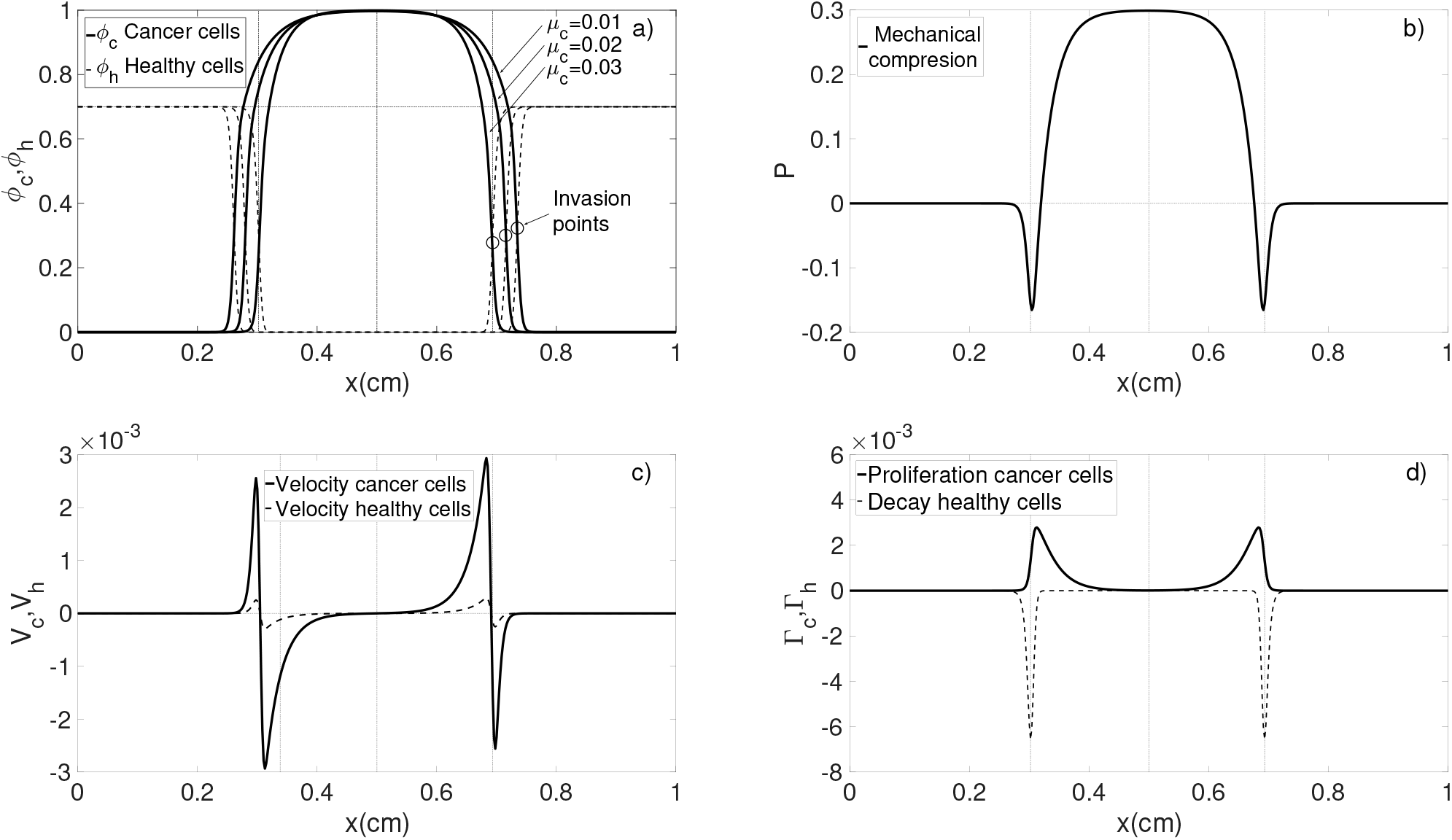
Physical properties of tumor growth at *t* = 8.7 months later without mass-effect. a) Cancer and healthy cell distribution profile with three different permeability coefficients. b) Mechanical pressure on the healthy tissue. c) The growth velocity of tumor and tissue. d) Proliferation and decay profile of cancer, and healthy cells.

The mechanical pressure is shown in Figure 4b). Based on the Fig. 4a) for *μ_c_* = 0.01, we observe that the mechanical load in the tumor region is proportional to the cancer cell density. The maximum compression is located in the tumor centre where the maximum cancer cell density occurs, and decreases gradually toward the edges as the tumor cell density becomes small. Notice that the mechanical load becomes negative in those regions where the population of cancer and healthy cells compete. The destruction of healthy cells by cancer cells leads to a void-effect with negative pressure, i.e., *ϕ_c_* + *ϕ_h_ < ϕ*_0_.

The corresponding velocity of cancer and healthy cells, obtained from Eqs. (11) and (12), are depicted in Fig. 4c). Comparing the profiles of the data with those of the Fig. 4a), the velocity of cancer cells is zero in the regions where the cancer cell density is highest. As the density decreases, cells are less packed so that they have more freedom of movement, and the velocity increases until it reaches its maximum value near the border where the cancer cells infiltrate the healthy tissue. The healthy cells begin to affect the velocity of the cancer cells, once the two populations come into contact. In other words; When *ϕ_c_ > ϕ_h_*, the velocity of cancer cells abruptly decreases (but remains positive) as cancer cells infiltrate the tissue. Such behavior is modified when *ϕ_h_ > ϕ_c_*. The resistance of the healthy tissue to being compressed and displaced pushes the cancer cells back, giving the cancer cells negative velocity. The profile of velocities of *V_c_* and *V_h_* are anti-symmetric around *x*=0.5. From Eqs. (7) and (8), we highlight that the amplitudes of the curves vary because of the different permeability coefficients of the cancer and healthy cells. In her model, Swanson reports that the velocity of cancer cells, (defined by 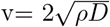 and determined via Fisher’s law,) is a constant with value v=0.008 cm/day. In our model, we find that the velocity of the cells depends on the population surrounding the cells, which have velocities that range from 0 to 0.002. This thus satisfies the condition 0 ≤v≤ 0.002 cm/day. Finally, in Fig. 4d), we analyze the curve of proliferation and decay of cancer, and healthy cells. We see clearly that the tumor is most active at its boundary, with increased growth rate for the cancer and increased decay rates for the healthy tissue.

Our second and more relevant study is performed when the anisotropy of the brain and the mass-effect - terms that play an important roles during the tumor progression- are simultaneously considered. The first term is replicated by an heterogeneous distribution of diffusion coefficients that represent the white fiber tracks in the brain, while the second one is the mechanical compression derived from competition for space between the growing tumor and the healthy tissue. In the next sections, we analyze the shape of the periphery of the tumor taking into account the anisotropic structure of the white matter using i) an artificial diffusion coefficient and ii) a more realistic structure provided by the Computer Tomography data.

### B. Anisotropy modeled by a linear diffusion coefficient

In order to gain experience about the effects of the anisotropy on the tumor shape, we study the progression of the lesion within a medium whose diffusion coefficient linearly grows from 0.0005 to 0.0020 cm^2^*/*sec, as depicted by the dashed line in Fig. 2. From the numerical simulation, depicted in Fig. 5, we observe that there is an small difference between the edges of the tumor that grows in an isotropic structure and that that grows in a linearly anisotropic structure. In general, we notice that the distance traveled by the cancer cells in the anisotropic case is less in the region *x <* 0.5 due to *D_c_* ⪅ 0.0013. Conversely, when *D_c_ >* 0.0013 just as it happens in *x >*0.5, the distance traveled is greater. We noticed, additionally that the response of the healthy tissue is proportional to the volume increased, as discussed before. In summarize, the lower (larger) the diffusion coefficient, the shorter (greater) the path traveled per unit of time by the cancer cells along the white matter, as noticed on the left (right) hand side of Fig. 5.

**Figure 5.**
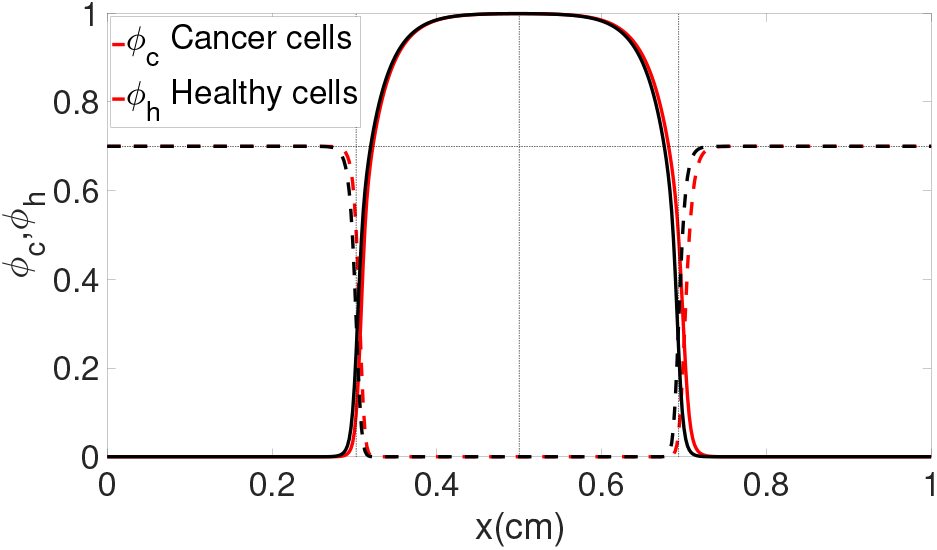
One-dimensional schematic representation of the distribution of cancer cells (solid) and healthy cells (dashed) of a virtual brain tumor. Black lines show the case of constant diffusion coefficient and red shows the spread using a linear increasing diffusion coefficient as shown in Figure 2. We see a small increase in the spread to the right, following the gradient of the diffusion coefficient.

### C. Anisotropy modeled via CT data

A more relevant and realistic study is performed when the anisotropy of the medium where the tumor reside is similar to the spatial structure of the white matter. For this aim, we consider that the initial stage of the virtual tumor is represented by a Gaussian function centered in *x*=0.5, and started numerically with the same parameters defined in Section A. The anisotropic architecture of the white and gray matter is described by the gray-scale image obtained from a Computer Tomography (CT), as depicted in Figure 6a) [62]. The CT image is a segment of brain, represented by a two-dimensional surface made up of pixels in gray-scale, whose numerical values range from 0 to 1 to distinguish the white from the gray matter in the brain. The method for artificial approximation of the diffusion coefficients in our 1-dimensional model consists of selecting a horizontal segment from the CT image, Figure 6b), and re-scaling their values to the order of the diffusion coefficients registered in the human brain, as depicted in Figure 6c). The artificial diffusion coefficients are used to evaluate the tumor progression numerically in time.

**Figure 6.**
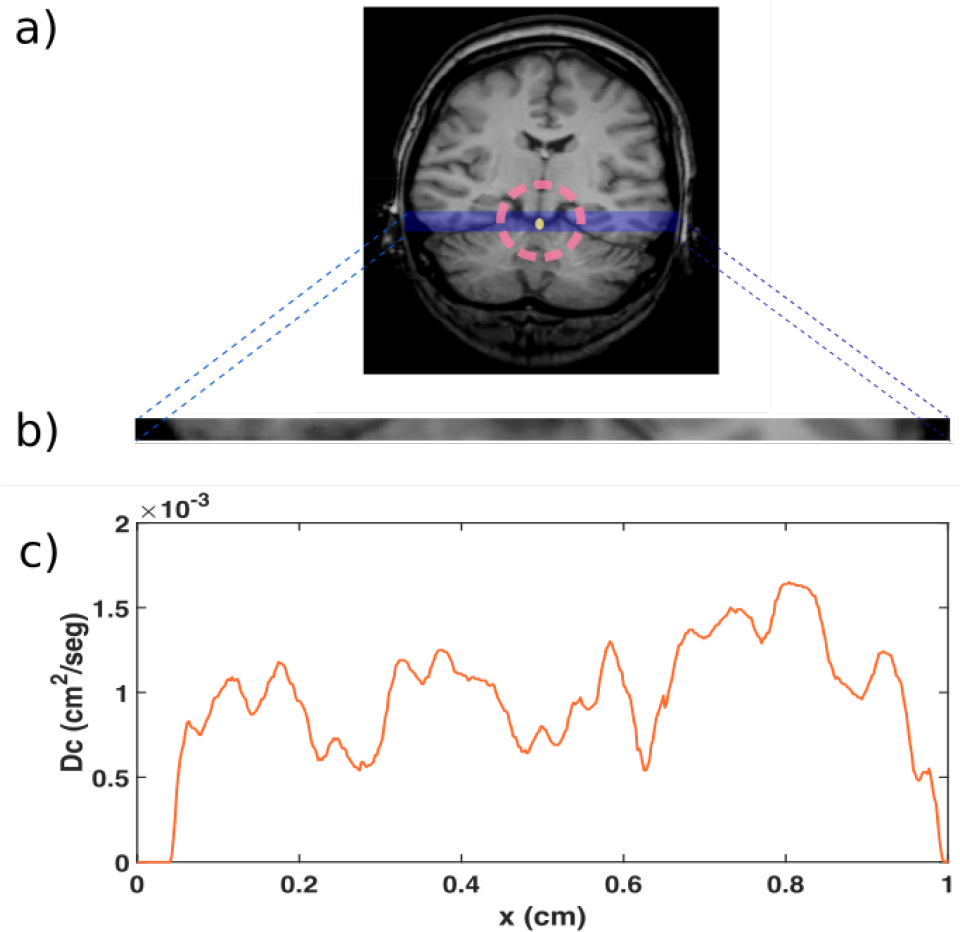
Virtual representation of the diffusion coefficients from a CT image. a) Slice of human brain obtained from CT. b) Segment of an CT. c) Artificial diffusion coefficients obtained from rescaled data of the CT.

The numerical solution to Eqs. (9) and (10) at *t* ≈8.7 months later is depicted in Fig. 7a). The image shows that the spatial anisotropy of the brain implemented via the diffusion coefficients modifies the growth patterns of the tumor during its spatio-temporal progression. In the figure, we observe that the numerical solution to our advection-diffusion model with the mass-effect allows visualizing an extended tumor that decays and infiltrates the healthy tissue anisotropically. The distribution of cancer cell density has its maximum density (red line) around *x*=0.5 and decreases asymmetrically as the cells spread out in the healthy tissue, reaching their lowest density once the cells have infiltrated the healthy tissue. We emphasize that the infiltration of the cells in the healthy tissue is asymmetric, and the diffusion coefficients play a central role in determining its extent. In general, we can say that the bigger the diffusion coefficient, the greater the population of cancer cells that diffuse in a particular direction and invade the healthy tissue. In this sense, our model satisfies the hypothesis that the cancer cells migrate in those directions that offer less resistance to the movement, suggested in [39]. The infiltration level in the healthy tissue can be visualized via the slope of the tangent lines touching the point *ϕ_c,h_*=0.5, on both sides of the curve (red line). On the right-hand side, the decay of cancer cells is slow but invasive, while on the left, the decay is fast but only mildly aggressive. The response of the healthy tissue to tumor growth, on the other hand, is different (blue line). Although the invasion by the cancer cells into the healthy tissue occurs at the same location, the response of the healthy tissue to being displaced and compressed by the cells depends on its local environment. The healthy tissue located at *x >* 0.5, has greater resistance to being displaced by the cancer cells, which happens on both sides of the tumor’s central axis at *x* = 0.5.

**Figure 7.**
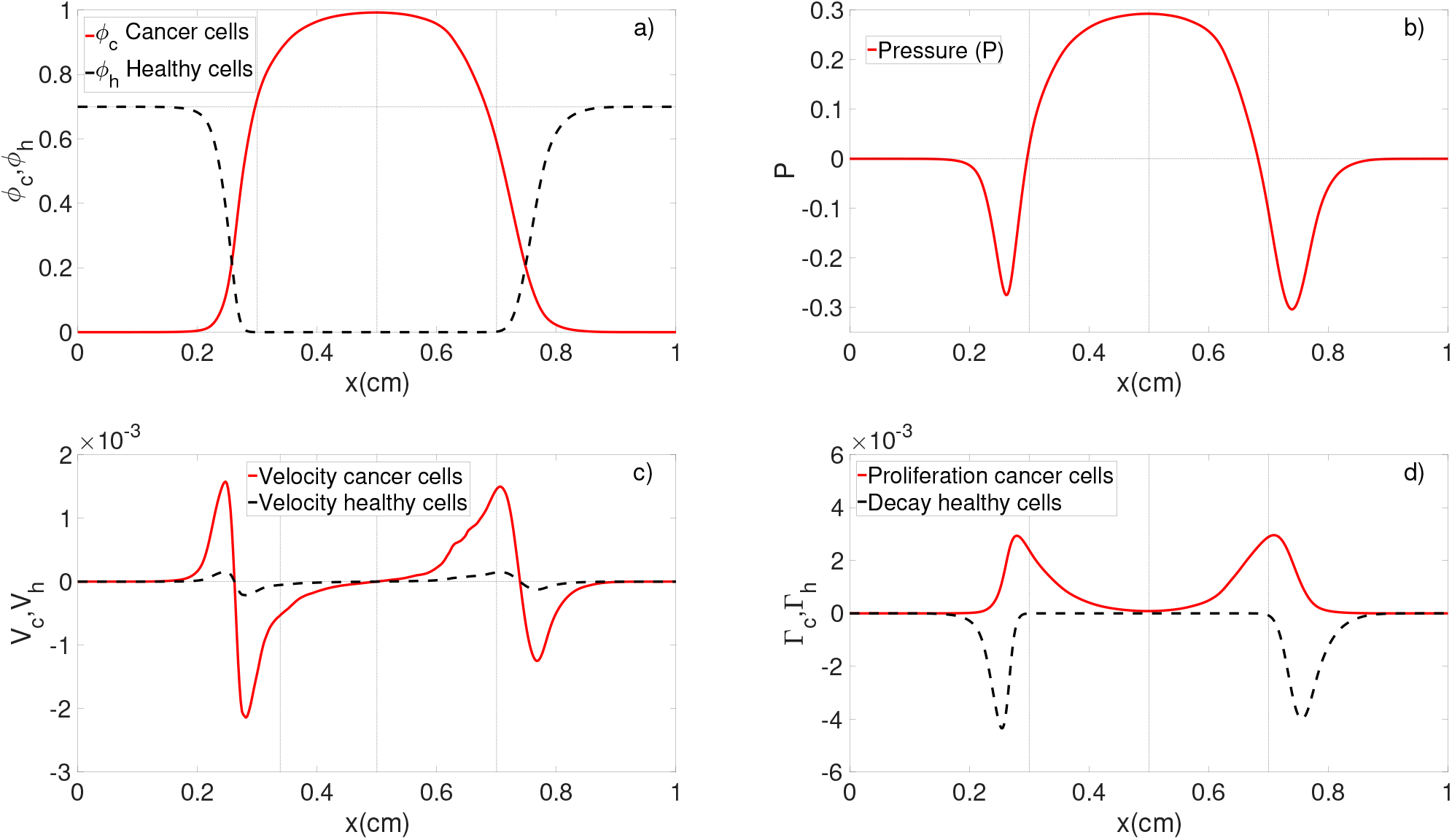
Physical properties of tumor growth at *t* ≈ 8.7 months with mass-effect. a) Cancer and healthy cell distribution profile with three different permeability coefficients. b) Mechanical pressure on the healthy tissue. c) The growth velocity of tumor and tissue. d) Proliferation and decay profile of cancer, and healthy cells.

The irregular growth, additionally, also affects the mechanical pressure between the cells. Referring to Fig. 7b), the pressure registered at each point of the tumor is directly proportional to the cell density in that location. The pressure is positive and asymmetric on both sides of *x*=0.5 and decreases until reaching negative values when *ϕ_c_ < ϕ*_0_ before reaching the tumor-brain interface. Notice that the lowest points of the curve are different. Their negative values are due to the density of cancer and healthy cells so that *ϕ*_0_ *> ϕ_c_* + *ϕ_h_*, and their different minimums refers to the different population density *ϕ_c_* + *ϕ_h_* in both sides.

As was discussed in Section A, the growth velocity of the tumor depends on the spatial distribution of both the cancer and healthy cells, as well as the anisotropic structure conferred to the white and gray matter. Making reference to Fig. 7c) (red line) and supported by 7a), the growth velocity has different behaviors: when *ϕ_c_ > ϕ_h_* = 0, the tumor growth velocity is exponential, as a consequence of the exponential proliferation. If *ϕ_c_ > ϕ_h_* ≠ 0, the exponential growth is interrupted abruptly by the presence of healthy cells in such a way that the growth velocity decays, reaching its null value when *ϕ_c_* = *ϕ_h_*. When *ϕ_c_ < ϕ_h_*, the cancer cells have infiltrated the healthy tissue so that the elevated concentration of healthy cells promotes negative velocities, whose values become zero as *ϕ_c_* approaches 0. Notice that the curve *V_c_* is asymmetric and the height of its peaks are not equal. The reason for this is that cancer, and healthy cells have different permeability coefficients that satisfy the inequality *μ_c_ > μ_h_*. This implies that the cancer cells have greater resistance to being invaded than do the cells of healthy tissue.

In the last Figure 7d), we display the profile of the proliferation and death rates of cancer and healthy cells, respectively. The cells exposed to elevated mechanical pressure have a low or null proliferative capacity, as depicted in Figure 7d). The apparent free proliferation of cancer cells is interrupted when cancer cells detect the presence of healthy cells that have invaded their space so that the proliferation starts to decay as the cancer cells sense the increasing population of healthy cells. The population of healthy cells, on the other hand, diminishes as the cancer cells invade the zone of the healthy tissue. This population decay is stopped at the visible interface between the tumor and the tissue. At this point, the decay of healthy cells gradually ceases until it is null. When the cells sense the maximum pressure, the proliferation rate is null, however, the proliferation rate increases as the pressure is reduced. The maximum proliferation rate occurs in those regions where the pressure is zero, but the rate decreases drastically all the way to zero, when the tumor reaches the zero-stress configuration. Conversely, as the tumor cells invade the healthy tissue, such cells decay gradually, reaching their minimum decay when the pressure is zero and increasing as the pressure increases again.

Finally, we describe the anisotropic proliferation of cancer and healthy cells. Referring to the Fig. 7d), we look again at the hypothesis that the proliferative capacity of the cancer cells of a tumor is inversely proportional to the cancer cell population density, and consequently to the pressure they perceive exerted upon themselves. The proliferation curve of cancer cells displayed in Fig. 7d) (red line) shows that the cells located in the nucleus of the tumor -where the cancer cell density is highest-have an extremely low proliferation rate. But this proliferative capacity changes depending on their locations. The closer the cells are to the nucleus, the lower their proliferation. However, such behavior is conditioned by the presence of healthy cells. We observe that if the glioma cells are isolated from healthy cells, their proliferation grows exponentially. And conversely, the proliferation decreases as the population of healthy cells increases.

## V. RESULTS AND CONCLUSIONS

The precise evaluation of the spatio-temporal infiltration of brain tumors remains a challenging task. Currently, the exact location of the tentacle-like branching of the tumor invading the healthy tissue cannot be defined with accuracy due to the lack of precision of the imaging devices, so that the life prognosis in patients is discouraging. Nowadays, the mathematical models play a central role in validating new proposals and comprehending underlying mechanisms of the tumor growth. In the vast scientific literature, we find that the proliferation, diffusion, and brain architecture, are the cornerstones of mathematical models of tumor growth. In recent times, the inclusion of mechanical aspects is being explored as a critical element to improve the predictive capacity of mathematical models. The central hypothesis of the mass-effect requires the assumption that the tumor growth occurs in a cavity with limited space and rigid boundaries. In this way, mechanical forces, derived from the lesion’s growth, affect the tumor shape and, hypothetically, regulate the direction of growth, which is guided towards regions of lower compression. One hypothesis suggests that such heterogeneous mechanical loads can induce apoptosis or cellular inhibition in regions of high compressive stress, and allow a proliferation of the low-stress areas of the tumor spheroids [4]. This mechanical analysis represents a significant contribution that can explain or predict tumor morphology. In this context, mechanical forces (or mass-effect) are being implemented as a guide element to improve the prediction of the tumor shape.

In the present work, we have proposed a one-dimensional mathematical model to explore the effects of mechanical pressure during the spatio-temporal evolution of a brain tumor. The implementation is modeled via the resultant mechanical force from the outward radial pressure generated from the tumor expansion, and the inward radial pressure caused by the resistance of the healthy tissue to being compressed. We identified that the implementation of the mass-effect during the tumor growth allows predicting a greater extension of the lesion. Our model suggests that the direction of more significant expansion (or tumor growth) depends on the population of cancer cells and mechanical pressure. The tumor model can be employed to predict the direction and extent of the spread of a tumor. Although this model lacks vital information from the patient for a fully adequate diagnosis, it does enable us to illustrate the impact that the mechanical compression, derived from the tumor’s growth, has upon its architecture.

## ACKNOWLEDGEMENT

I am grateful for helpful discussions and guidance by Dr. T. Hillen and to the University of Alberta for its support. I thank CONACyT-Mexico for the financial support during my Postdoctoral Research.

## References

[1] George Paxinos Charles Watson, Matthew Kirkcaldie. The Brain: An Introduction to Functional Neuroanatomy. Elsevier, 2010.

[2] L. Angela Mihao. Silvia Budday, Gerhard A. Holzapfel, Ellen Kuhl, and Alain Goriely. A family of hyperelastic models for human brain tissue. J. of the Mechanics and Physics of Solid, 106:60–79, 2017.

[3] Emad Moeendarbary and Andrew Harris. Cell mechanics: Principles, practices, and prospects. Wiley Interdisciplinary Reviews: Systems Biology and Medicine, 6:371–88, 09 2014.

[4] Rakesh K. Jain, J. D. Martin, and T. Stylianopoulus. The role of mechanical forces in tumor growth and therapy. Annu. Rev. Biomed. Eng., 16:321–346, 2014.

[5] Kolahi KS and Mofrad MR. Mechanotransduction: a major regulator of homeostasis and development. WIREs Syst. Biol. Med., 2:625–639, 2010.

[6] Alf Gerisch Tommaso Lorenzu Chiara Villa, Mark A. J. Chaplain. Mechanical models of patterns and form in biological tissues: the role of stress-strain constitutive equations. arXiv, (q-bio.TO), 2021.

[7] Alain Goriely, Marc G. D. Geers, Gerhard A. Holzapfel, Jayaratnam Jayamohan, Antoine Jérusalem, Sivabal Sivaloganathan, Waney Squier, Johannes A. W. van Dommelen, Sarah Waters, and Ellen Kuhl. Mechanics of the brain: perspectives, challenges, and opportunities. Biomech Model Mechanobiol, 14:931–965, 2015.

[8] Weaver VM. Barnes JM, Przybyla L. Tissue mechanics regulate brain development, homeostasis and disease.. J Cell Sci., 30(1):71–82, 2017.

[9] C. Nelson and M. Bissell. Of extracellular matrix, scaffolds, and signaling: tissue architecture regulates development, homeostasis, and cancer. Ann. Rev. Cell Dev. Biol., 22:287–309, 2006.

[10] J. Lotem and L. Sachs. Epigenetics and plasticity of differentiation in normal and cancer stem cells. Oncogene, 25(59):7663–7672, 2006.

[11] Lam PY, Di Tomaso E, Ng HK, Pang JC, Roussel MF, and Hjelm NM. Expression of p19ink4d, cdk4, cdk6 in glioblastoma multiforme. Br. J. Neurosurg, 14:28–32, 2000.

[12] Friedman A. Bigner D. et al. McLendon, R. Comprehensive genomic characterization defines human glioblastoma genes and core pathways. Nature, 445:1061–1068, 2008.

[13] Nayak A, Ralte AM, Sharma MC, Singh VP, Mahapatra AKand Mehta VS, and Sarkar C. p53 protein alterations in adult astrocytic tumors and oligodendrogliomas. Neurol India., 52:228–32, 2004.

[14] Stevens B. Salter, M. Microglia emerge as central players in brain disease. Nat Med, 23:1018–1027, 2017.

[15] Chia-Ching John Lin, Kwanha Yu, Asante Hatcher, Teng-Wei Huang, Hyun Kyoung Lee, Jeffrey Carlson, Matthew C Weston, Fengju Chen, Yiqun Zhang, Wenyi Zhu, Carrie A Mohila, Nabil Ahmed, Akash J Patel, Benjamin R Arenkiel, Jeffrey L Noebels, Chad J Creighton, and Benjamin Deneen. Identification of diverse astrocyte populations and their malignant analogs. Nature Neuroscience, 20(3):396–405, 2017.

[16] B. Alberts, J. D. Watson, J. Lewis, D. Bray, K. Roberts, and M. Raff. Molecullar Biology of the cells, 4th edition. New York: Garland Science, 2002.

[17] J. F. Holland, E. Frei, R. C. Bast, D. W. Kufe, D. L. Morton, and R. R. Weichselbaum. Cancer Medicine, 3rd ed. Lea a Febiger, Philadelphia, 1993.

[18] Vincent T. DeVita, Steven A. Rosenberg, and Theodore S. Lawrence. Cancer: Principles and Practice of Oncology, 9th ed. Lippincott, Philadelphia, 2011.

[19] Reifenberger G von Deimling A Figarella-Branger D Cavenee WK Ohgaki H Wiestler OD Kleihues P Ellison D. Louis DN, Perry A. The 2016 world health organization classification of tumors of the central nervous system: a summary. Acta Neuropathol, 131(6):803–820, 2016.

[20] H. J. Scherer. Structural development in gliomas. American Association for Cancer Research Journals, 34(3):333–351, 1938.

[21] Murray J.D. Mathematical Biology, 3rd ed. New York, NY: Springer-Verlag, 2002.

[22] Antonio Omuro and Lisa M. DeAngelis. Glioblastoma and Other Malignant Gliomas: A Clinical Review. JAMA, 310(17):1842–1850, 11 2013.

[23] Urbanska K., Soko-lowska J., Szmidt M., and Sysa P. Glioblastoma multiforme-an overview. Contemp Oncol (Pozn)., 18(5):307–12, 2014.

[24] K.R. Swanson, Jr E.C. Alvord, and J.D. Murray. A quantitative model for differential motility of gliomas in grey and white matter. Cell Prolif., 33:317–329, 2000.

[25] Anne Line Stensjøen, Ole Solheim, Kjell Arne Kvistad, Asta K. Håberg, Øyvind Salvesen, and Erik Magnus Berntsen. Growth dynamics of untreated glioblastomas in vivo. Neuro-Oncology, 17(10):1402–1411, 2015.

[26] Woodard D.E. Bartoo G.T. Murray J.D.-Alvord Jr E.C. Tracqui P., Cruywagen G.C. A mathematical model of glioma growth: the effect of chemotherapy on spatio-temporal growth. Cell Prolif., 28:17–31, 1995.

[27] Patricia K. Burgess, Paul M. Kulesa, James D. Murray, and Ellsworth C. Alvord Jr. The Interaction of Growth Rates and Diffusion Coefficients in a Three-dimensional Mathematical Model of Gliomas. J. of Neuropathology and Experimental Neurology, 56(6):704–713, 1997.

[28] Larry A. Taber. Biomechanics of Growth, Remodeling, and Morphogenesis. Applied Mechanics Reviews, 48(8):487–545, 1995.

[29] KR Swanson, EC Alvord Jr., and JD Murray. Virtual brain tumours (gliomas) enhance the reality of medical-imaging and highlight inadequacies of current therapy. British Journal of Cancer, 86:14–18, 2002.

[30] Kristin Swanson. Virtual and real brain tumors: using mathematical modeling to quantify glioma growth and invasion. J. Neurol. Sci., 216(1):1–10, 2003.

[31] Chaplain M.A. and Sleeman B.D. A mathematical model for the growth and classification of a solid tumor: a new approach via nonlinear elasticity theory using strain-energy functions. Math. Biosci., 2:169–215, 1992.

[32] Macklin P. and Lowengrub J. Nonlinear simulation of the effect of microenvironment on tumor growth. J. Theor Biol., 245(4):677–704, 2007.

[33] J. M. Maloney, D. Nikova, F. Lautenschl ′’a ger, E. Clarke, R. Langer, J. Guck, and K. J. Van Vliet. Mesenchymal stem cell mechanics from the attached to the suspended state. Biophysical journal, 99(8):2479–2487, 2010.

[34] Villa-Diaz LG Sun Y, Lam RHW, Chen W, Krebsbach PH, and Fu J. Mechanics regulates fate decisions of human embryonic stem cells. PLoS ONE, 7(5):e37178, 2012.

[35] K. Pogoda, L. Chin, P. C. Georges, F. J. By-field, R. Bucki, R. Kim, M. Weaver, R. G. Wells, C. Marcinkiewicz, and P. A. Janmey. Compression stiffening of brain and its effect on mechanosensing by glioma cells. New journal of physics, 16:075002, 2014.

[36] Baker A M, Bird D, Lang G, Cox T R, and Erler J T. . New journal of physics, 32:1863–8, 2013.

[37] Samuel M S et al.. Cancer Cell, 19:776–91, 2011.

[38] Olsen A L, Bloomer S A, Chan E P, Gaca M D, Georges P C, Sackey B, Uemura M, Janmey P A, and Wells R G. Am. J. Physiol. Gastrointest Liver Physiol., 301:G110–8, 2011.

[39] Stylianopoulos T., Martin J.D., Chauhan V.P., Jain S.R., and et al. Diop-Frimpong B. Causes, consequences, and remedies for growth-induced solid stress in murine and human tumors. Proc. Natl. Acad. Sci. USA, 109:15101–8, 2012.

[40] Tse J Cheng G and Munn LL Jain RK. Micro-environmental mechanical stress controls tumor spheroid size and morphology by suppressing proliferation and inducing apoptosis in cancer cells. PLoS ONE, 4(4):e4632, 2009.

[41] D. Ambrosi and F. Guana. Mechanics aspects of growth in soft tissues. Boll. Unione Mat. Ital, 7:775–781, 2004.

[42] D. Ambrosi and F. Guana. Strees-modulated growth.. Math. Mech. Solids, 12:319–343, 2007.

[43] A. Jones, H. Byrne, J. Gibson, and J. Dold. Mathematical models for the stress induced during avasculas tumor growth. J. Math. Biol, 40:473–499, 2000.

[44] R. Araujo and D. McElwain. A linear-elastic model of anisotropic tumor growth. Eur. J. Appl. Math., 15:365–384, 2004.

[45] G. Vitale L. Preziosi. Mechanical aspects of tumour growth: Multiphase modelling, adhesion, and evolving natural configurations. New Trends in the Physics and Mechanics of Biological Systems, Lecture Notes of the Les Houches Summer School, Oxford University Press, 92:177–228, 2011.

[46] H. Byrne and L. Preziosi. Modelling solid tumor growth using the theory of mixtures. Mathematical Medicine and Biology, 2001(7):547–581, 2001.

[47] L. Preziosi and A. Farina. On Darcy’s Law for Growing Porous Media. International Journal of Non-Linear Mechanics, 37(3):485–491, 2002.

[48] D. Ambrosi and F. Mollica. The role of stress in the growth of a multicell spheroid. Journal of Mathematical Biology, 48(5):477–499, 2004.

[49] Burton A.C. Rate of growth of solid tumours as a problem of diffusion. Growth, 30:157, 1966.

[50] Greenspan H. P. Models for the growth of a solid tumor by diffusion. Studies in Applied Mathematics, 51:317, 1972.

[51] Thomlinson R. H. and L. H. Gray. The histological structure of some human lung cancers and the possible implications for radiotherapy. Br. J. Cancer, 9(4):539–49, 1955.

[52] Greenspan H. P. On the Growth and Stability of Cells cultures and Solid Tumors. J. theor. Biol., 56:229–242, 1976.

[53] Olivier Clatz, Pierre-Yves Bondiau, Hervé Delingette, Maxime Sermesant, and Simon K. Warfield et al. Brain Tumor Growth Simulation. [Research Report] INRIA, RR-5187:1–45, 2004.

[54] Davide Ambrosi and Luigi Preziosi. Cell adhesion mechanisms and stress relaxation in the mechanics of tumours. Biomech Model Mechanobiol, 8:397–413, 2009.

[55] E. De Angelis and L. Preziosi. Advection-Diffusion models for solid tumor evolution in vivo and related free Boundary problem. Math. models and Methodos in Appl. Sciences, 10(3):379–4907, 2000.

[56] A. Swan, T. Hillen, J. Bowman, and A. Murtha. An anisotropic model for glioma spread. Bulletin Math. Biol., 80, 2017.

[57] K.J. Painter and T. Hillen. Mathematical modeling of glioma growth: The use of diffusion tensor imaging data to predict the anisotropic pathway of cancer invasion. Journal of Theoretical Biology, 323:25–39, 2013.

[58] A patient-specific in vivo tumor model. Mathematical Biosciences, 136(2):111–140, 1996.

[59] J. P. Ward and J. R. King. Mathematical modelling of avascular-tumour growth. IMA J. Math. Appl. Med. Biol., 14:39–69, 1997.

[60] J. P. Ward and J. R. King. Mathematical modelling of avascular-tumour growth II: Modeling growth saturation. IMA J. Math. Appl. Med. Biol., 15:1–42, 1998.

[61] J. P. Ward and J. R. King. Mathematical modelling of the effects of mitotic inhibitors on avascular tumour growth. J. Theor. Med., 1:171–211, 1999.

[62] https://community.wolfram.com/groups/-/m/t/294122.

